# Introducing CHiDO – a No Code Genomic Prediction Software implementation for the Characterization & Integration of Driven Omics

**DOI:** 10.1101/2024.03.05.583604

**Authors:** Francisco González, Julián García-Abadillo, Diego Jarquín

**Affiliations:** Agronomy Department, University of Florida, Gainesville, FL 32611; Centro de Biotecnología y Genómica de Plantas, Universidad Politécnica de Madrid (UPM), Campus de Montegancedo, Pozuelo de Alarcón, Madrid, 28223, Spain

**Keywords:** Multi-Omics Integration, Genomic Selection - Genomic Prediction, R Shiny, Climate Adaptation, Low-code-no-code (LCNC), Bayesian Statistics, High Dimensional Interactions

## Abstract

Climate change represents a significant challenge to global food security by altering environmental conditions critical to crop growth. Plant breeders can play a key role in mitigating these challenges by developing more resilient crop varieties; however, these efforts require significant investments in resources and time. In response, it is imperative to use current technologies that assimilate large biological and environmental datasets into predictive models to accelerate the research, development, and release of new improved varieties. Leveraging large and diverse data sets can improve the characterization of phenotypic responses due to environmental stimuli and genomic pulses. A better characterization of these signals holds the potential to enhance our ability to predict trait performance under changes in weather and/or soil conditions with high precision. This paper introduces CHiDO, an easy-to-use, no-code platform designed to integrate diverse omics datasets and effectively model their interactions. With its flexibility to integrate and process data sets, CHiDO’s intuitive interface allows users to explore historical data, formulate hypotheses, and optimize data collection strategies for future scenarios. The platform’s mission emphasizes global accessibility, democratizing statistical solutions for situations where professional ability in data processing and data analysis is not available.

**Core ideas:**

1. The authors developed CHiDO, a platform for breeders to build predictive models integrating multi-omics data.
2. CHiDO is a no-code tool that leverages the reaction norm model proposed by Jarquin et al. (2014).
3. The platform aims to increase access to predictive analytics lowering relevant technical and financial barriers.

## 1 INTRODUCTION

As the global population continues to surge, projected to reach 10 billion by 2050 (Gu et al., 2021), the imperative to increase agricultural yields becomes increasingly critical (Van Dijk et al., 2021). This challenge is compounded by the escalating frequency and intensity of weather variations due to climate change, posing a significant threat to food security worldwide (Lesk et al., 2016). Such climatic extremes have already begun to impact the productivity of elite crop varieties, with studies indicating a potential reduction of up to 6% for an increase of one degree Celsius in average temperature (Zhao et al., 2017).

To meet these rising demands for available food products, agricultural production must increase, and supply chain improvements must be achieved to reduce food waste at different stages. Plant breeding can play a pivotal role in increasing total harvestable output through the development of improved genotypes that can withstand changing environmental conditions.

More resilient crop varieties serve dual purposes: *1*) fulfilling the nutritional demands of a growing population, and *2*) mitigating reliance on environmentally harmful inputs like fossil fuels and synthetic agrochemicals (Foley et al., 2011). However, traditional plant breeding methods, which predominantly rely on phenotypic and pedigree data, are resource-intensive and time-consuming (Atlin et al., 2017). These conventional approaches require significant investments in land and time, often taking up to eight years to develop a new variety for annual crops (Jarquín et al., 2017). Moreover, genetic engineering, while a potential solution, is surrounded by socio-economic and ecological concerns, as well as issues of accessibility, corporate control and public acceptance (Clapp, 2018; Tsatsakis et al., 2017).

Recent advances in sequencing technologies have revolutionized our ability to characterize genotypes with high precision through DNA-based marker profiles (Varshney et al., 2014). This genomic information enables the characterization of genetic relationships between individuals (Bernardo 1994), forming the foundation of genomic selection (GS). However, this selection framework has its own set of challenges, such as handling large genomic datasets for reduced number of phenotypic observations --the “large *p*, small *n*” problem--. Leveraging genomic selection-by-genomic prediction (GS-GP) techniques allows the prediction of the performance of unobserved genotypes based solely on their marker profiles (Meuwissen et al., 2001). This approach, although a significant leap in breeding efficiency, overlooks the impact of environmental factors. The next iteration of computational methods to accelerate and improve breeding efforts was modeling Genotype-by-Environment (G×E) interactions.

G×E analysis examines how genotypes respond to different environmental conditions (change in the response patterns - rankings). However, similar problems (*p>>n*) than for conventional GS-GP models arise when considering the interaction between genes and environmental factors increasing the computational demands of modeling a large number of interactions via contrasts (Crossa et al., 2017). Utilizing the approach proposed by Jarquín et al. (2014), we can overcome these challenges by analyzing and integrating all first degree interactions between marker SNPs and environmental covariates via covariance matrices/structures. This alternative significantly reduces the dimensionality of data by leveraging correlations between genotype-by-environment combinations that are similar both genetically and at the level of the environmental covariates (environmentally) rather than computing individual contrast effects between markers and weather factors (Jarquín et al., 2014). Several studies have shown the advantages of taking into consideration the G×E interaction in prediction models in plant and animal breeding applications (Jarquin et al., 2020, Tiezzi et al., 2017). The predictive power of the G×E interaction can be bolstered through the inclusion of a broad spectrum of omics (or layers) data (e.g. genomics, proteomic, metabolomics, enviromics, ionomics, high-throughput data, etc.), known as multi-omics analysis (Yang et al., 2021).

Implementing models that effectively integrate and interpret this complex multi-omics data can be challenging, often requiring specialized programming and statistical expertise that may not be readily available in many breeding programs around the world, especially in developing regions. To address this gap, we have developed CHiDO, a no-code platform designed to facilitate the integration of multi-omics data to build, train and test complex G×E prediction scenarios.

Across several Latin-American cultures, the word ‘chido’ (meaning ‘cool’ in English) is a powerful and oversimplistic expression that succinctly describes all the positive aspects of an action, event, thing, etc. In our case, CHiDO stands for **Ch**aracterization and **I**ntegration of **D**riven **O**mics, and it enables breeders to use advanced analytical methods without having to write code themselves; removing a technical barrier and democratizing access to the latest predictive analytics used in breeding implementations. Our CHiDO development is not just a “prediction software”, it also integrates a series of developments proposed by several Latin American scientists (Drs. de los Campos, Crossa, Perez-Rodriguez, Gianola) that are recurrently cited along this paper, and this is a way to acknowledge their contributions in the field. In this paper, we discuss the development of this platform, its components and the statistical methods leveraged for its functionality. Currently, the application can be accessed at https://jarquinlab.shinyapps.io/chido/.

## 2 MATERIALS AND METHODS

### 2.1 Platform Overview

The implementation of elaborated prediction models, integrating data from multiple omics of information (including interactions of several types), and their corresponding evaluation considering different prediction scenarios (mimicking realistic scenarios) poses extra challenges in many breeding programs.

ChiDO is a no-code platform that can fill this gap by allowing breeders to build, train, and validate linear models that incorporate data derived from multiple omics of information in a simple manner. It also addresses two challenges with leveraging G×E interaction models for breeding efforts by *1*) using a UI-based workflow to overcome the technical barrier associated with multi-omics data handling and programming, and *2*) reducing the dimensionality of G×E by adopting the reaction norm model described in Jarquín et al., (2014) which is further described in the *Statistical Background* subsection. The platform’s user interface (UI) is divided into four sections –data loading, model assembly, training/validation, and results view where each section contains instructions and widgets to customize the metadata, parameters and model equations as necessary. The drag-and-drop interface is a novel approach to building complex models where users can add individual omics as main effects and form interactions between them (e.g., G×E) by *collapsing* these effects without requiring any advanced programming knowledge.

Other key features of CHiDO include: *1*) customizable data processing and parameter tuning, *2*) handling multiple input files within a single session, *3*) viewing omics data and editing associated metadata, *4*) building and testing multiple models in a single session, *5*) viewing results in the UI with the option to download them as CSV and PNG files, and *6*) exporting models as R objects via an RDS file. These features and additional functionality are split into four separate page views within the CHiDO platform (**Figure 1**).

**Figure 1.**
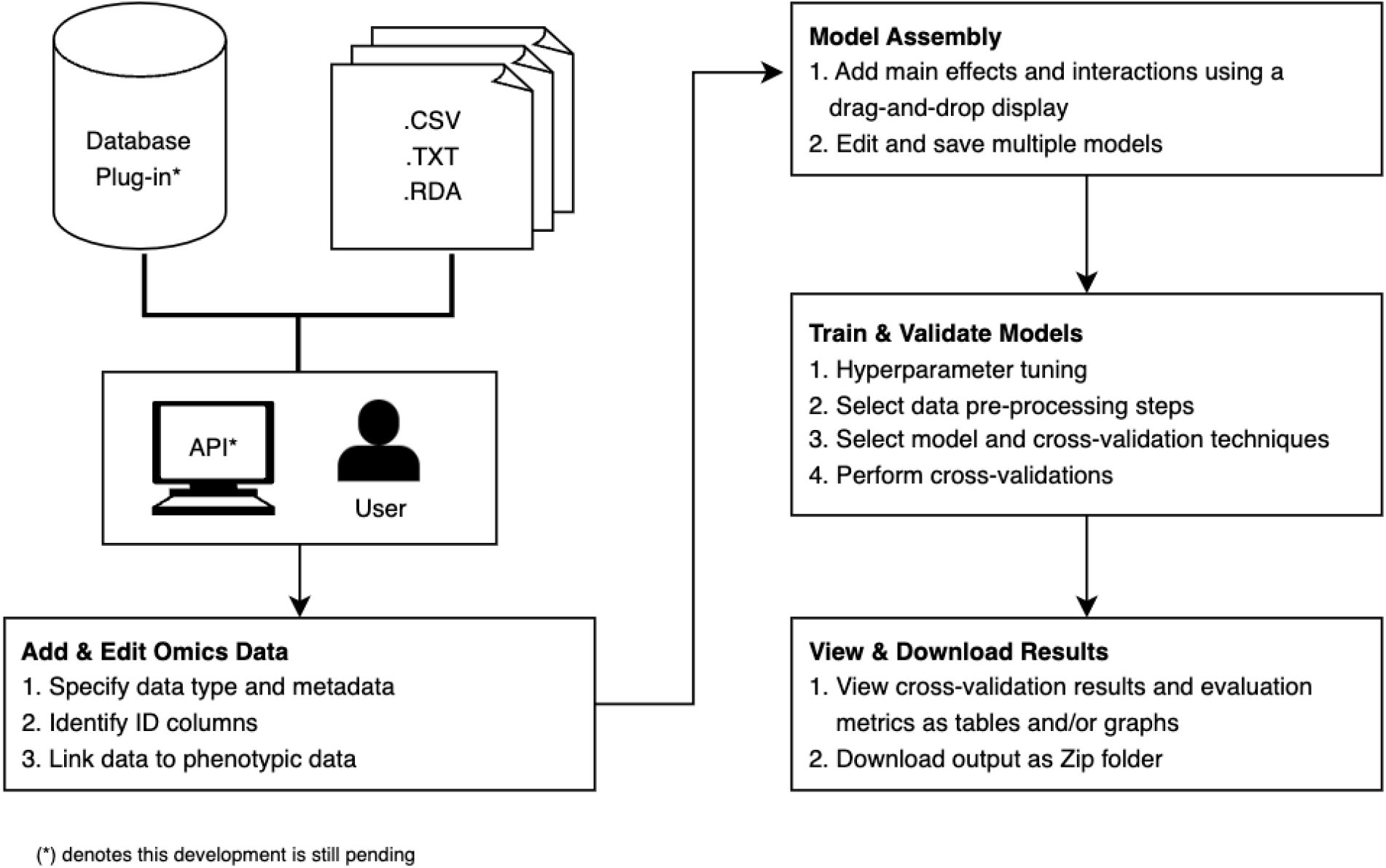
Overview of the different components and functionalities within the CHiDo platform

#### 2.1.1 Design

CHiDO was built using the *Shiny* framework (Winston Chang et al., 2023), a popular R package for creating interactive web and desktop applications. Its architecture, however, diverges from the typical UI-Server split typically found in most *Shiny* applications, emphasizing a modular design methodology. This approach involves segmenting logical components into individual functions to enhance the platform’s long-term maintainability, support and expansion. The software’s architecture is based on modern development practices to prevent logical duplication, reduce dependencies within the codebase, and minimize disruption as new versions are released.

Consequently, CHiDO’s design integrates both R and JavaScript in its frontend and backend logic. The packages *shinyjs* (Dean Attali, 2021) and *shinyjqui* (Yang Tang, 2022) are utilized to introduce functionality that extends beyond the traditional capabilities of R/Shiny, including the drag-and-drop interface for model assembly (Figure 2). Many features are made available by leveraging a suite of R packages such as *ggplot2* (Hadley Wickham, 2016) for rich data visualizations and *DT* (Yihui Xie et al., 2023) to display and handle tabular objects. The selection of *DT* is deliberate, enabling table and data frame manipulation with either JavaScript or R code. For an exhaustive list of libraries used by CHiDO, see Table 1.

**Table 1.** List of all R packages used in the creation of CHiDO.

| Package | Function in CHiDO | URL |
| --- | --- | --- |
| shiny | Main framework for displaying and organizing the web application | https://CRAN.R-project.org/package=shiny |
| shinydashboard | Simplified the creation of dashboards within the Shiny framework | https://CRAN.R-project.org/package=shinydashboard |
| gridExtra | Align widgets, plots, and data in grid-like format | https://CRAN.R-project.org/package=gridExtra |
| dplyr | Perform data processing and transformations in a consistent manner | https://CRAN.R-project.org/package=dplyr |
| DT | Handle and render tabular objects using R and/or JavaScript syntax | https://CRAN.R-project.org/package=DT |
| ggplot2 | Generate graphics of cross-validation results and evaluation metrics | https://ggplot2.tidyverse.org |
| shinyjs | Integrating JavaScript into Shiny application to extend functionalities of UI | https://CRAN.R-project.org/package=shinyjs |
| shinyjqui | Enable animation effects needed for the drag-and-drop interface in the model assembly page | https://CRAN.R-project.org/package=shinyjqui |

**Figure 2.**
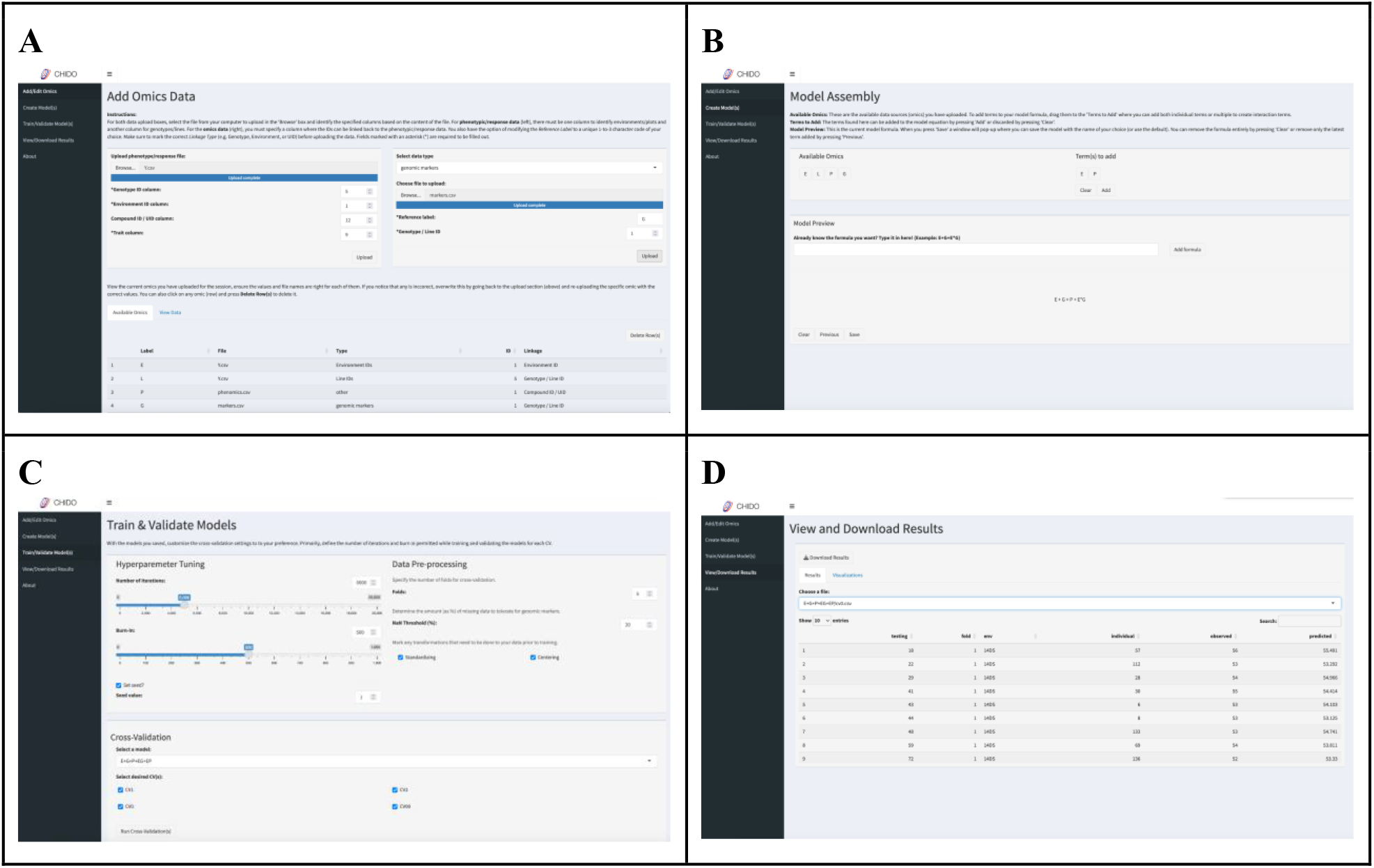
User interface for CHiDO; (A) The *Add Omics Data* page is where users upload their files and define metadata for each of them such that the platform can treat them as separate omics; (B) The *Model Assembly* page lets users create multiple models using the uploaded data as main effects or combining them with interaction terms; (C) Users can tune training and validation parameters, apply quality control on the genomic data, as well as selecting the different cross-validation schemes to employ; and (D) the *View and Download Results* page allows users to view prediction outputs and evaluation metrics in tabular and graphical formats before downloading them to the user’s local environment.

#### 2.1.2 Usage

The typical workflow for CHiDO (Figure 3) can be listed as the following steps:

**Figure 3.**
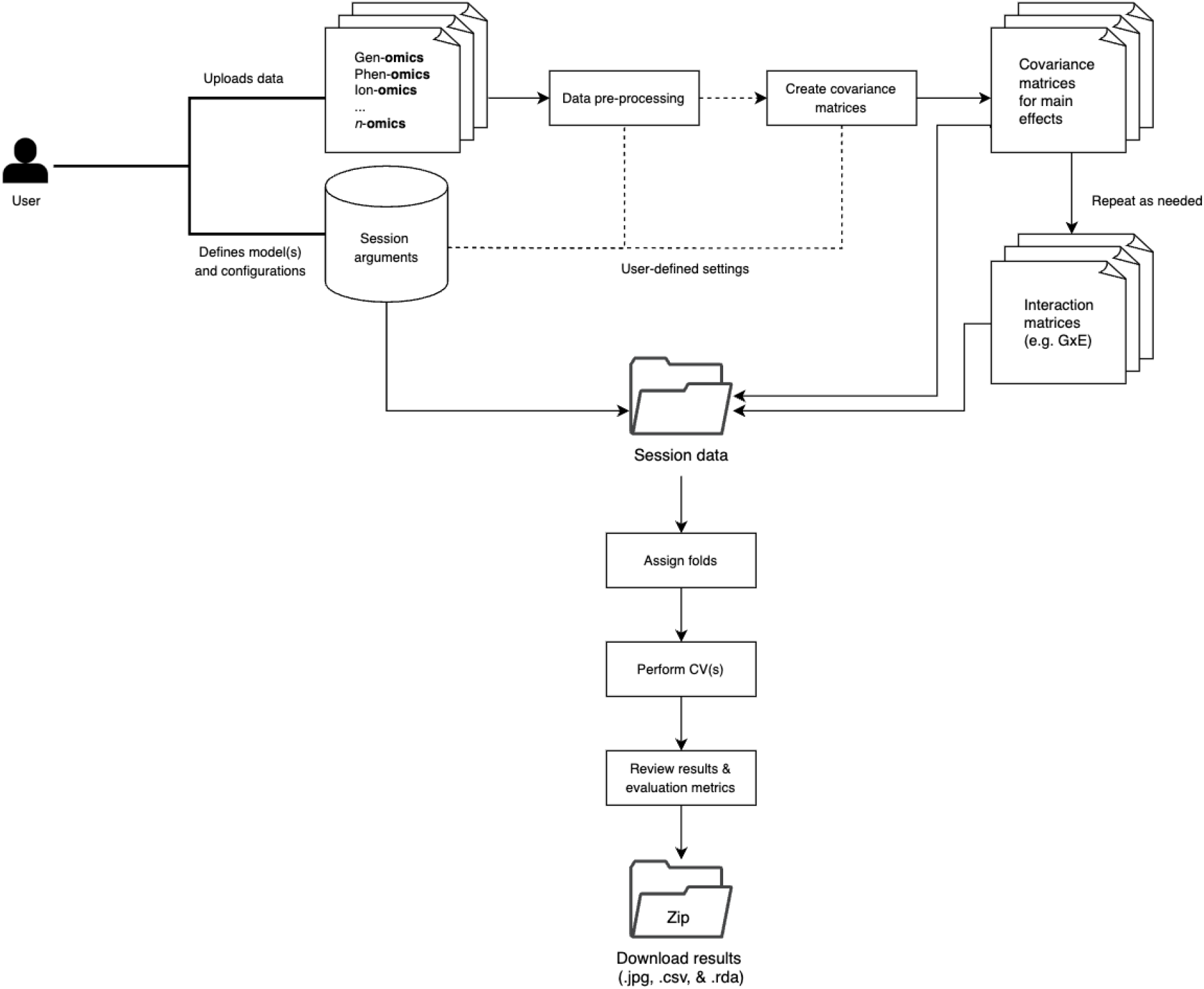
Workflow diagram for CHiDO. This diagram demonstrates the logic implemented in the application to create and train linear models using arguments and data provided by the user.

**Step 1:** CHiDO accommodates the upload of tabular data in CSV and RDA format, with a flexible approach to data requirements. The primary necessity is the phenotypic response file (Y), which must contain columns for environment IDs, genotype IDs, and the target trait to predict, at minimum. Dealing with data from multi-environment trials, the column corresponding to the ID of the environments, and the genotypes should be specified in the interface. Also, if omic data is collected at the plant or sample level (e.g. multispectral data collected with drones, ionomic data, information collected on secondary traits, etc.) a column serving as a unique identifier (*compound*) for that level should be included for alignment.

For omics data, each file must have an identifier column specified by the user to link back to the matrix of phenotypes in the Y file; this could be a column with the genotype IDs, an environment ID column, or a unique identifier (UID-*compound*) column (e.g. genotype-in-environment combination, plant or sample ID). Once a data file is uploaded, users can modify its metadata including its display label and linkage type (Environment ID, Genotype / Line ID, Compound ID).

Users are responsible for ensuring their data is properly formatted prior to uploading it to CHiDO. Extra care should be taken to ensure that all identifier levels of an omic are represented in the Y file. For example, if molecular information is uploaded, all lines referenced in this dataset should be present in the Y file as part of the Genotype / Line ID column, even if the corresponding phenotypic values are missing. If these identifiers are not consistent across both files, the covariances matrices cannot be constructed for the implementation to work.

**Step 2:** Upon upload, each file is recognized as a separate omic within CHiDO and is assigned a unique label, if not specified by the user during upload. In the model assembly section, these labels appear as draggable elements for the user to add as main effects into a linear model formula box. Interaction effects can be added by dragging -*collapsing*-two or more of these labels into the same box before adding them into the formula. Users can build and save as many models as desired, facilitating comparative analysis of these to determine which set of effects can best predict trait performance (e.g., including G×E interactions) for desired phenotypic expression.

**Step 3:** In the training and validation section, users have the option to adjust convergence hyper-parameters (e.g., number of iterations and burn-in) and data pre-processing steps on genomic data (i.e., quality control – minor allele frequency and percentage of missing values) at their discretion. These settings can be altered for each model or applied uniformly across the multiple models created in the previous section. Once a model is selected, it can be tested with one or more of the four distinct cross-validation (CV) schemes that mimic prediction scenarios of interests to breeders; *1*) CV2: predicting tested genotypes in observed environments; *2*) CV1: predicting untested genotypes in observed environments; *3*) CV0: predicting tested genotypes in new environments; and *4*) CV00: predicting untested genotypes in new environments (Persa et al. 2021). The implementation of the declared linear predictors (models) is done using the BGLR (Bayesian Generalized Linear Regression) package developed by Perez and de los Campos (2014).

**Step 4:** The results of the selected CV schemes are presented in the UI in both tabular and graphical outputs, with the option to download these locally. The downloadable results are delivered in a compressed zip folder where the contents are systematically sorted by model, with each model’s folder containing CSV files with the raw numeric data for each CV and PNG files showing a graphical representation of the results. In addition to the CV data, the results also include evaluation metrics to assist with model interpretation efforts and the corresponding variance components derived from the full data analysis. The metrics available are *prediction accuracy* (as the Pearson correlation between predicted and observed values), *root-mean-squared-error* (RMSE), and *variance components* to evaluate the relative contribution of each one of the omics to explain the phenotypic variability. The formulae for each metric are provided in the *Statistical methods* section. The output and evaluation metrics are displayed in both tabular and graphical formats. This data is available to view as overall model performance (Figure 4) or split by environment (Figure 5).

**Figure 4.**
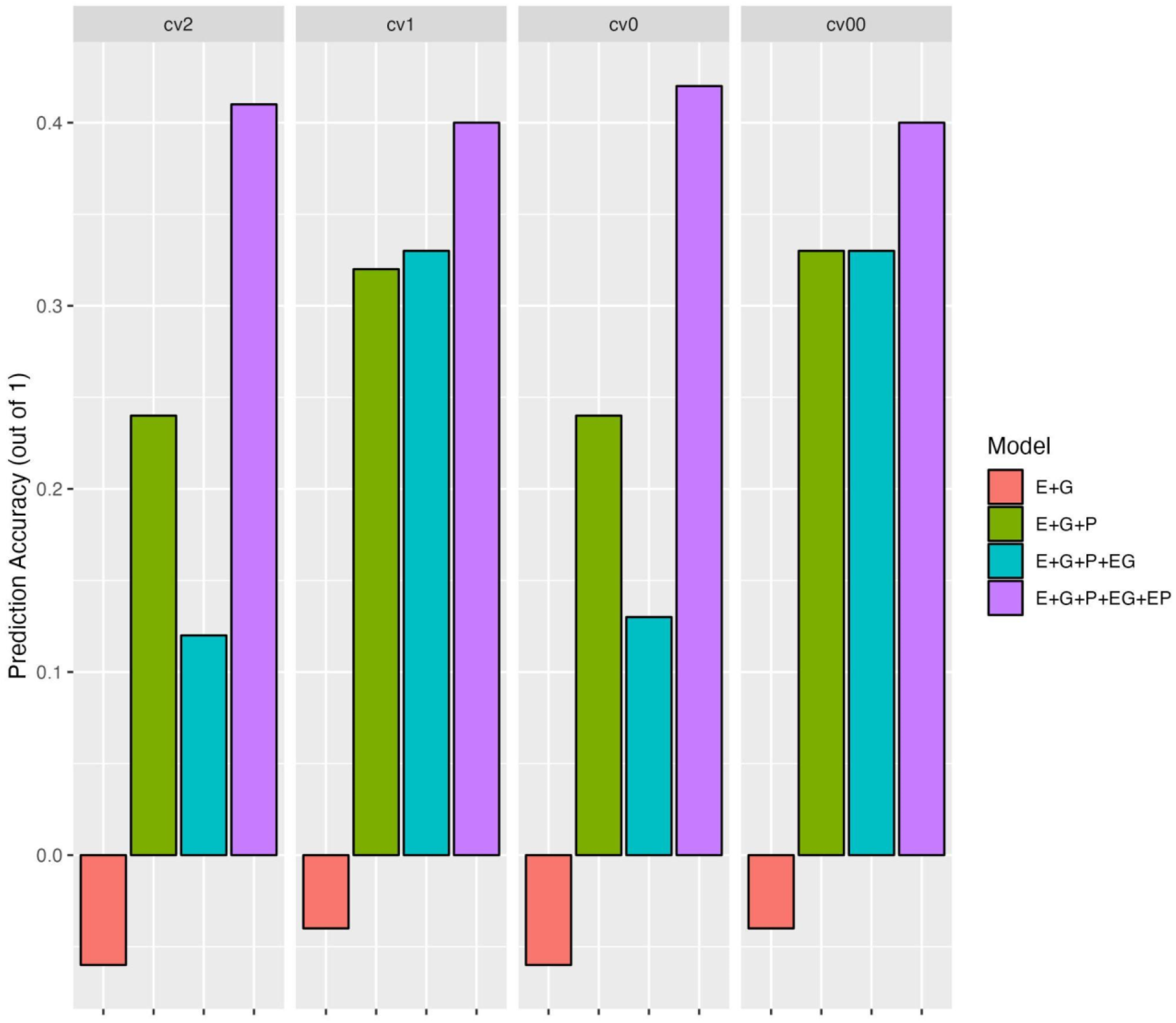
Example of prediction accuracy results by model, and by cross-validation scheme.

**Figure 5.**
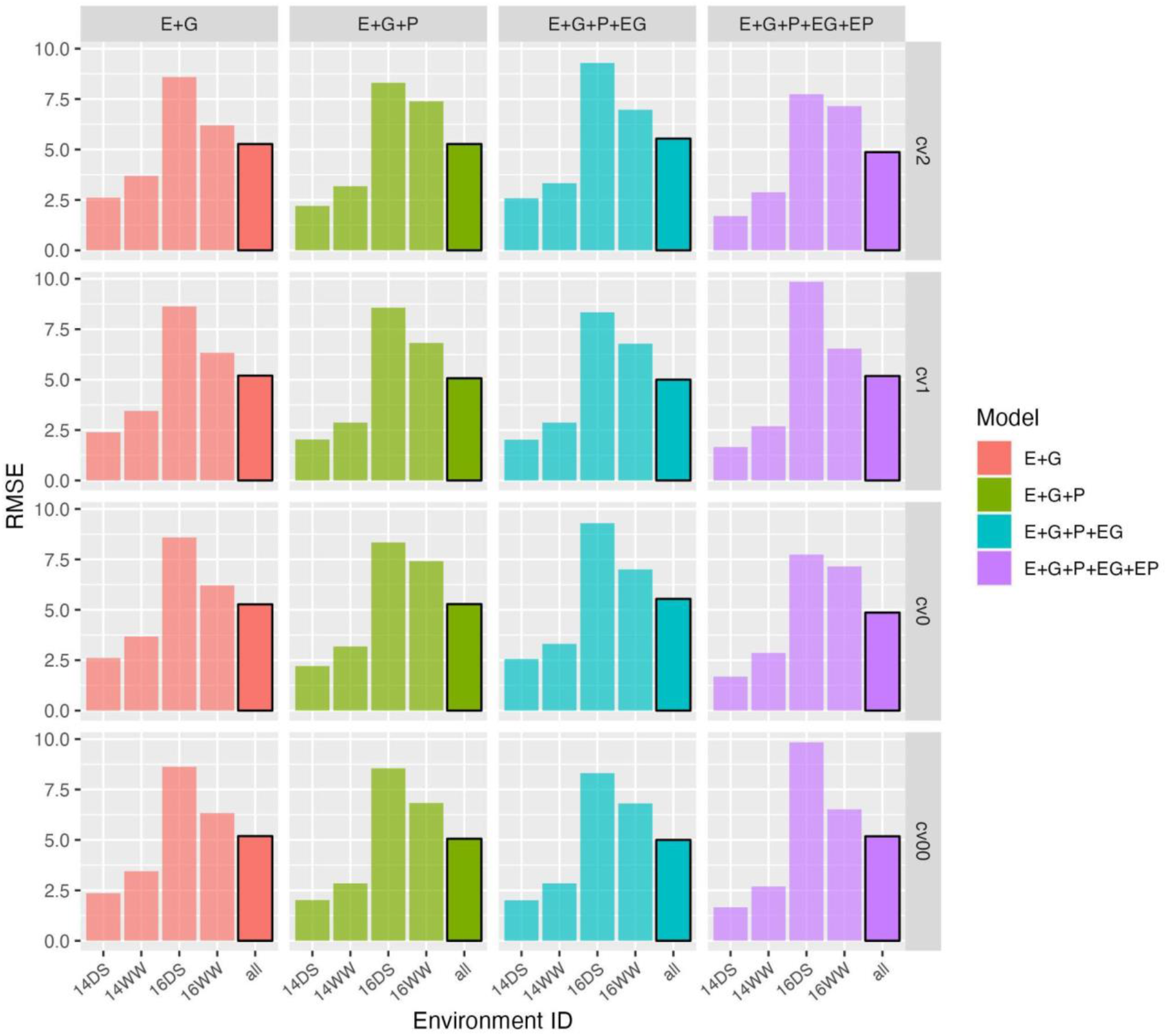
Example of the model root-mean-square error (RMSE) by environment, and by cross-validation scheme.

### 2.2 Statistical Background

Modeling high-dimensional sets of covariates *p* (e.g., genomic, environmental, gene × environment interactions, etc.) using a reduced set of *n* phenotypic observations such that *p >> n*, poses extra challenges. Especially under the conventional prediction approaches based on linear regressions of the ordinary least squares (OLS) framework. The phenotypic response *y*_*i*_ of the *i*^th^ genotype (*i* = 1, 2, …, *n*) can be represented as the linear combination between *p* markers *x*_*ij*_ (*j* = 1, 2, …, *p*) and their corresponding effects *b*_*j*_ such that

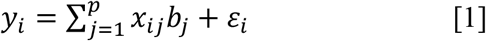

Then, under the OLS framework the solution for the vector of marker effects is given by

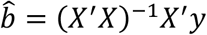

A major challenge to obtain the solution of the vector of marker effects is the inversion of singular matrices of the form (*XX*^′^)^−1^ which are not full rank due to the larger number of coefficients to estimate (*p*) with respect to the reduced number of data points (*n*) available for model fitting. Under the parametric context, several statistical approaches have been developed to deal with the course of the dimensionality (*p >> n*). Two of the most popular statistical frameworks are the penalized regressions and the Bayesian methods which in many cases are a sort of Bayesian versions of the former ones. By design, the penalized methods delimit to *n* the total number features or covariates to select in the model.

On the other hand, the Bayesian methods consider distributional assumptions of the marker effects, allowing (in principle) all features to be included in the final prediction model. In both cases, the inversion of matrices with large dimensions (*p × p*) is accomplished by adding a value to the diagonal elements of the (*XX*^′^) matrix to “*break*” the singularity. Another option is to consider prior distributions for the marker effects such that 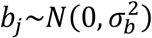. This will help to reduce the uncertainty of their estimation (*prediction*) by adding a bias. In both cases, the general solution takes the following form

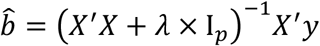

The value to add in the diagonal matrix of *X*^′^*X* is conveniently selected in a trade-off between model goodness of fit and model complexity, where 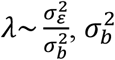, and 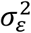 are the corresponding variance components of the effect of the genomic covariates (genes/SNPs) and of the error term.

Although the previous implementations allow us to get a solution when considering main effects only, these still deal with large matrices and do not solve the problem of including interactions between high dimensional sets of covariates (e.g., *p* genomic or phenomic and *Q* environmental features for a total of *p* × *Q* first order contrasts). To tackle this problem, first we examine an alternative parameterization proposed in animal breeding (VanRaden, 2008) to include main effects in a computationally-convenient manner, then we provide a few details of the implemented method for including interactions between groups of covariables.

The Genomic Best Linear Predictor (G-BLUP) attempts to directly compute the genomic effect of the *i*^th^ individual g_*i*_ resulting from the linear combination between *p* marker and their corresponding effects such that 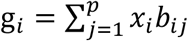. Hence, instead of focusing in obtaining first the marker effects *b*_*ij*_ to be used later in the linear combination, the genomic effect is obtained in one step. The solution to this model requires the inversion of matrices of the type (*XX*^′^)^−1^ and order *n* × *n* instead of *p* × *p*, facilitating the handling of information derived from large matrices, with *X* centered and scaled by columns (rows-genotypes; columns-marker SNPs). Under this parameterization, the vector of genetic effects is modeled as 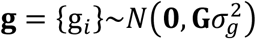, where 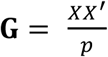 and 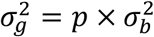. Here, **G** corresponds to the kinship matrix whose entries describe the genomic similarities between pairs of individuals (VanRaden 2008). The resulting model in a matrix parameterization is as follows

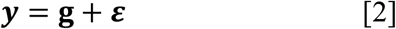

A similar idea based on covariance structures can be considered to include high-dimensional interactions between factors (more details are provided below in 2.3 section). Jarquin et al. (2014) proposed the reaction norm model that allows the inclusion of all first order interactions between genomic and weather factors. First, it was shown that the main effects of weather covariables can be introduced into models in a similar fashion than the main effect of genes or marker SNPs. Here, the environmental similarities between pairs of environments can be characterized using weather information. This is analogous to considering marker SNPs to conducting the genomic characterization between pairs of genotypes. Then the interactions between markers and environmental factors are introduced via covariance structures computed as the element-to-element product between the previous covariance matrices for genotypes and environments.

Indirectly, this model, below described, allows to include the interaction between each marker SNP and each weather covariate by modeling the interaction between their corresponding linear combinations via covariances structures following the G-BLUP model fashion. The resulting covariance structure of this interaction component that considers genomic and weather factors is computed as the Hadamard product, which is the cell-by-cell product between two or more covariance structures of the same dimension. In this case, the corresponding covariance structures are redistributed/extended according to the vector of phenotypes and levels of the corresponding factors (genotypes and environments) to ensure these are conformable.

In summary, modeling the G×E interactions can be both computationally and statistically expensive due to the high dimensionality of the number of contrasts that can be formed between genetic markers and environmental covariates (ECs). There are equivalent methods that reduce such dimensionality by introducing markers and ECs via covariance structures as described in Crossa et al. (2017). Interactions can be introduced through covariance structures computed via the Hadamard product between these. Although these methods were already developed, there is no simple method for capturing and integrating interactions among different omics.

### 2.3 Statistical Methods – Model Building

As mentioned previously, CHiDO was developed as a way to easily build models that can capture main effects of diverse omics and incorporate the interactions between these, such as those derived between genomic markers and ECs. CHiDO’S drag-and-drop interface simplifies the process of creating complex models and adds a layer of abstraction for the methodology established by Jarquin et al. (2014).

#### 2.3.1 Main effects

Upon uploading the phenotypic response file, CHiDO automatically recognizes the environment (*E*) and genotypic line (*L*) data to make them available as random effects, ***E***_*j*_ and ***L***_*i*_, respectively. These random effects can be added as terms in the model assembly section to capture the inherent variability in phenotypic responses due to environmental and genetic differences. Therefore, a base model with no additional omics data can be represented as

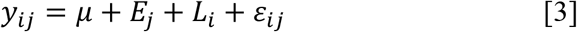

where *y*_*ij*_ is the phenotypic observation (target trait) of the *i*^th^ genotype (*i* = 1, …, *L*) in the *j*^th^ environment (*j* = 1, …, *J*), *μ*is the overall mean and *ε*_*ij*_ is the error term capturing the non-explained variability by the other model terms.

When an omic data set **O** is uploaded, CHiDO attempts to compute its specific vector of effects **o** = {o_*k*_}, transforming the data into a covariance matrix **Ω** that captures the similarities among the pairs of entries for the different factor values (e.g., genotypes, environments, genotypes-in-environments, etc.) For instance, if a file containing *p* (*m* = 1, …, *p*) genetic markers ***X*** = {*x*_*im*_} is uploaded, CHiDO attempts to modeling the vector of genomic effects g = {g_*i*_} as described for the G-BLUP model by constructing a genomic relationship matrix **G** whose entries describe genomic relationships between pairs of genotypes. For a given factor *f* (e.g., genotype, environment, genotype-environment combination) with *T* levels (*t* = 1,…, *T*), the generalized form of the vector of effects associated to an omic-type **o** can be calculated as a linear combination between *M* covariates *O*_*tm*_ and their corresponding effects *τ*_*m*_ (e.g., SNP markers, weather covariates, soil features, multispectral, Near InfraRed NIR, etc.)

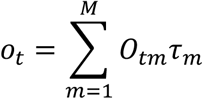

Using this form, we can describe general main effects for a given factor. For example, modeling the genomic effect of the *i*^*th*^ (*i* = 1, 2, …, *L*) genotype using marker information *X* = {*x*_*il*_} on *p* molecular markers and their corresponding effects *b*_*l*_ (*l* = 1, 2, …, *p*) we have

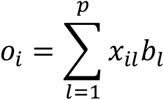

Similarly, for modeling the effect of the *j*^*th*^ environment based on *Q* weather covariates *W* = {*W*_*jl*_} and their corresponding effects *γ*_*l*_ (*l* = 1, 2, …, *Q*) we have

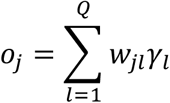

For an omic data observed at the particular/specific level -*compound-* (e.g. genotype-in-environment combinations; the *i*^*th*^ genotype in the *j*^*th*^ environment), such as those derived from high-throughput phenotyping platforms, the information *Z* = {*z*_*ijl*_} on *s* features (e.g., images) can be modeled also as a linear combination considering their corresponding effects *δ*_*l*_ (*l* = 1, 2,…, *s*) as follows

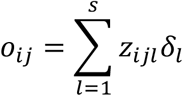

In addition, the information of covariance structures relevant to the factors of study (e.g., genotype, environment, etc.), can be also included in the models. For example, genetic effects based on the pedigree matrix **A**, or the environmental effects based on an environmental kinship matrix **C**. In these cases, it is necessary to specify the factor ID in the phenotypic matrix to connect with the associated covariance structure. Hence, the alignment of the data will be conducted as previously described, and also similar distributional assumptions (normality) as before will be considered such that

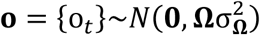

where **Ω** is the corresponding covariance structure whose entries describe similarities between pairs of levels (genotypes, environments, genotype-in-environments, etc.), and 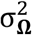 is the associated variance component. In this case, **Ω** might represent the pedigree matrix (**A**) whose entries describe genetic similarities between pairs of individuals. Also, **Ω** can represent an environmental kinship matrix (**C**) whose entries describe environmental similarities between pairs or environments. If a covariance structure derived from soil information (**S**) is available, it can be also introduced into the models in a similar manner.

Models including only main effects can be easily constructed by adding the information of the different omics into the linear predictor. For example, a linear model created using two omics, one generic of type **o** with *T*-levels (*t* = 1, 2, …, *T*) and *M* covariates, and another based on *p* genetic markers for *L* individuals (*i* = 1, …, *L*) can be represented by

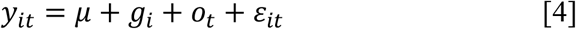

CHiDO’s capacity to handle different omic types of variable dimension extends to the creation and alignment of distinct covariance matrices for each associated dataset. This is done by calculating the matrix cross-product that reflects/expands the specific relationships across the levels of that omic type according to the matrix phenotypic responses. For this, on each of the different *F* omics **O**_*f*_ (*f* = 1, 2, …, *F*) it is necessary to compute the incidence matrix Z_*f*_ that connects phenotypes with the *T* different levels of the omics (e.g., genotype, family, environment, mega-environment, farm, herd, genotype-in-environment combination, etc.) Then, the resulting aforementioned covariance matrices are aligned and/or expanded across all phenotypic records by computing 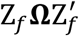 with the tcrossprod(tcrossprod(Z_*f*_,**Ω**), Z_*f*_) instruction.

#### 2.3.2 Multiplicative interactions

Interactions between different omics are modeled by calculating the Hadamard product of their corresponding covariance structures. For example, the interaction between the covariance matrices **G** and **Ω** denoted by (**G # Ω**), represents the interaction between genotypes (using molecular marker information **X)** and any other related omic -**O-**. The corresponding covariance matrix of this interaction term is represented with 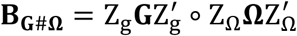, and modeled as

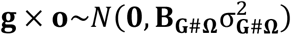

where Z_g_ and Z_Ω_are the corresponding incidence matrices that connect the phenotypic observations with the different levels of the omics (e.g., genotypes, environments, genotype-in-environment, families, etc.)

For any given covariance matrix of the main and interaction effects, CHiDO performs the spectral decomposition using the eigen() function to retrieve its *eigenvalues* and *eigenvectors*.

The *eigenvalues* reveal the magnitude of variance in the omic data along the directions defined by their corresponding eigenvectors. For G×E predictions, this information could provide insights into the major factors that contribute towards trait variation. This factorization is conveniently implemented to save computing time when fitting different linear predictors and prediction scenarios in BGLR R-package. Each time the BGLR function is used, and the covariance matrices are provided, it internally computes the eigen-value decomposition before starting the model fitting. Using datasets with a large number of phenotypic observations (*n*) this procedure might be time consuming, especially in those cases where the cross-validations involve exhaustive scenarios and/or folds. Thus, by providing the resulting factorization of these matrices a considerable amount of time and resources are saved avoiding extra computational burden.

#### 2.3.3 Cross-validation schemes

Prior to implementing prediction models in real-world applications such as GS, it is necessary to evaluate their usefulness integrating different omics to deliver accurate and reliable results.

Cross-validation studies are a common, time-tested method to perform such evaluation. Hence, after the models are created and saved in CHiDO, users can select from a range of cross-validation (CV) schemes (based on their specific research objectives) how to train and evaluate the performance of their model(s).

These CV schemes mimic real life prediction problems that breeders face at different stages along the breeding pipeline for the development of improved genotypes. As discussed, CV1 considers the prediction of ‘newly’ or untested genotypes in environments where other genotypes were already observed. CV2 (or incomplete field trials) mimics the prediction of already tested genotypes observed in other environments but not in the target environment (where other genotypes were also already tested). CV0 (or forward prediction) emulates the prediction of already tested (in other environments) genotypes in novel environments where no phenotypic records on any of the lines have been collected. CV00 is similar to the previous scheme with the main difference that the genotypes to predict have not been observed at any of the environments in the training sets. This last prediction scenario is the most challenging and probably the most interesting for breeders.

The manner to create the different partitions representing training and testing sets depends on the prediction problem (cross-validation scheme). Here, the folds are defined by the user according to the selected CV scheme to partition the phenotypic data (training/testing). For instance, in a *k*-fold cross-validation setting such as in CV1 and CV2, the dataset *D* is divided into *k* mutually exclusive subsets (*D*_1_, *D*_2_, …, *D*_*k*_), with each subset serving as a testing set -one at a time-while the remaining subsets are aggregated to form the training set. Under CV2 scheme, the phenotypes are randomly assigned to the folds, while under the CV1 scheme extra care is taken to assign genotypes to folds ensuring that all the phenotypic records from the same individual appear in the same fold. On the other hand, under CV0 and CV00 each environment naturally becomes a fold and care is taken to ensure similar training sample sizes to those in the previous schemes (CV2 and CV1) according to Persa et al., (2021). When performing the different CV schemes, CHiDO loops the folds until all folds are considered as testing or prediction sets using the BGLR function.

Since the models are fitted under the Bayesian framework, the users can define additional training hyper-parameters for BGLR such as the number of iterations and the burn-in rate. These parameters influence the convergence and stability of the Bayesian models. As mentioned above, the cross-validations are executed using the BGLR() function, which applies the user-defined settings. The *eigenvalues* and *eigenvectors* for each omic-matrix, carrying the information of the different model terms, are incorporated into the ETA object to compose the readable linear predictor for BGLR.

#### 2.3.4 Metrics

Upon completion of the BGLR analysis, CHiDO employs the model outputs to calculate several metrics essential for evaluating the performance of the different linear models. Custom functions have been developed within the CHiDO framework to facilitate these calculations, ensuring accuracy and efficiency in metric derivation.

##### Prediction accuracy (PA) measured on a trial basis

It is obtained by computing the Pearson’s moment correlation *ρ* between predicted and observed (phenotypic) values within each trial/environment/year/location/etc. This metric helps to determine how well a given model can predict phenotypic traits based on the multi-omics data associated to the provided model terms.

The formula for PA in the *j*^th^ environment (or grouping factor) is given by

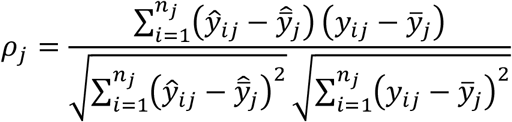

where 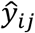 and *y*_*ij*_ are the predicted and the observed values of the *i*^th^ genotype at the *j*^th^ environment, 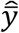 and 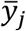 are their corresponding means, and *n*_*j*_ represents the number of observations at the *j*^th^ environment.

For an easier assessment of the model’s performance across environments, the weighted mean correlation is computed accounting for the uncertainty and the sample size of the environments according to Tiezzi et al. (2017) as follows:

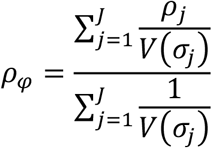

where 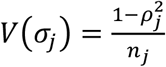 corresponds to the sampling variance.

##### Root Mean Squared Error (RMSE)

Quantifies the average magnitude of prediction error, measures a model’s precision, and penalizes large errors to a greater extent by squaring the difference between predicted and observed values. The formula for RMSE for the *j*^th^ environment is given by:

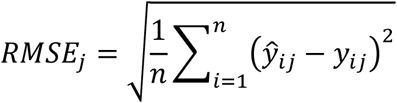

##### Variance Components

This metric measures the portion of variance explained by each model term associated to an omic with respect to the overall phenotypic variability. This estimation is critical for understanding which main or interaction effects influence the most the phenotypic expression/variability of target traits. It is computed considering a full data analysis (i.e., no missing values are generated on the phenotypic information).

The variance component of each term is computed as the percentage of the total variance explained, which for the *f*^th^ (*f=*1, 2, …, *F*) omic **O**_*f*_ it corresponds to the ratio between the current variance component and the sum of all the *F* variance components plus the unexplained residual variance 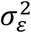

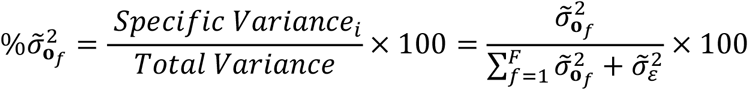

Here, under a given model, the specific variance refers to the variance attributable to the particular model term *f*^th^ and total variance is the sum of variances of all terms, including the residual variance. The total variance corresponds to the 100% of the phenotypic variability.

## 3 RESULTS AND DISCUSSION

Dealing with prediction analyses for breeding applications, usually an important amount of time (∼85%) is dedicated to the data preparation (quality control, alignment, cross-validation scenarios, etc.) and the remaining time (∼15%) is for the development and implementation of these models. Therefore, the availability of low-code, no-code (LCNC) applications such as CHiDO can help breeders save time and obtain expedited results by automating and assisting with many of these tasks, allowing them to focus on specific research questions derived from initial quick analyses.

In this paper we discussed the reason for CHiDO’s development, the technical and statistical methods applied, and its potential benefits to breeders. The CHiDO platform is a significant contribution to empower breeders and democratize access to modern solutions by enabling the modeling of different interaction types such as the G×E interaction without the need for in-depth programming knowledge. Increasing access to advanced analytics and prediction tools can not only accelerate research for new improved varieties/individuals, but also enable broader participation in agricultural research. CHiDO reflects a growing trend towards more accessible and flexible computational tools in genomics, as evidenced by recent literature advocating for the democratization of data science (Shang et al., 2019).

The practical implications of the CHiDO platform extend significantly beyond the immediate sphere of plant breeding. By enabling more accurate and efficient selection processes, CHiDO contributes to the development of crops with improved yields and environmental resilience. This capacity is particularly crucial in the context of climate change and the increasing demands for sustainable agricultural practices. The forthcoming introduction of interactive graphics for model evaluation further underscores CHiDO’s potential to enhance understanding and application of complex genomic data in breeding strategies.

LCNC platforms such as CHiDO are becoming increasingly popular and offer various benefits for researchers (Sufi, 2023). Some benefits include *1*) ease of adoption through a reduced learning curve, *2*) accelerated development speed, and *3*) circumventing resource scarcity, among many others listed in (Sufi, 2023; Yan, 2021). Despite these benefits, LCNC solutions are not without their challenges. Some notable drawbacks to LCNC are recurring costs and vendor lock-in. Similarly, developers can learn how to use the platform effectively but are bound to the limitations of said platform without the potential to extend its functionalities as opposed to custom developed alternatives.

We are addressing these drawbacks in CHiDO by ensuring the platform remains a free-to-use service and providing users the ability to submit issues or product feature requests on GitHub (https://github.com/jarquinlab/CHiDO). In addition to this, we are evaluating the potential release of CHiDO’s backend logic as an R package or API for more advanced users to extend CHiDO’s functionalities or integrate the tools with other packages when scripting.

In addition to the aforementioned features, future updates to CHiDO aim to enhance its functionality to cover a broader array of plant and animal breeding prediction scenarios, with the potential to extend these to public health applications such as personalized medicine. However, working on these proposed developments below detailed while maintaining an optimum functionality of the software will require of the investment of resources. We will seek for funding opportunities and partnerships to secure the needed resources to continue these and other future developments.

A few of the key additions we would like to integrate are modules for sparse testing designs, estimation of G×E markers using weather data -enabling a focused analysis on the relevance of genetic markers and ECs influencing target traits-, hybrid prediction via general and specific combining ability (GCA, SCA) terms and their corresponding interactions. Separately, CHiDO will incorporate options for selecting from multiple artificial intelligence (AI/ML) algorithms to facilitate the modeling of complex, non-linear relationships within multi-omics datasets. The use of Deep Learning and ML algorithms (e.g., RandomForest) is already being evaluated for their robustness in capturing intricate G×E interactions (Crossa et al., 2019), potentially leading to more accurate genomic selections. The launch of CHiDO online, alongside comprehensive documentation, is poised to democratize access to these advanced tools, stimulating worldwide collaboration and further research.

Ultimately, CHiDO stands at the forefront of integrating multi-omics data for plant breeding, representing a critical advancement in computational tools within agriculture. Its development is timely, addressing the urgent need for innovative solutions in plant breeding to meet the global challenges of food security and sustainability.

## ACKNOWLEDMENTS

JGA would like thank to the ‘Programa Propio’ of the ‘Universidad Politécnica de Madrid’ (UPM) for the financial support during his PhD studies and internship at the University of Florida. All authors would like to thank the UF Strawberry Breeding Program for their contribution in the technical discussion during the design phase.

## AUTHOR CONTRIBUTIONS

**FG** Methodology, Software, Validation, Formal analysis, Writing – Original draft, Writing – Reviewing and Editing; **JGA** Software, Visualization, Writing – Reviewing and Editing; **DJ** Conceptualization, Resources, Supervision, Writing – Reviewing and Editing

## DATA AVAILABILITY

There are no original data associated with this article. CHiDO is a web-based application accessible at https:jarquinlab.shinyapps.io/chido/ where users can upload their own data to develop predictive models. Data uploaded to CHiDO is not stored anywhere and is only used during the active session while users interact with the platform.

For demos and testing purposes, users can use sample data available at https://github.com/jarquinlab/CHiDO. These data sets were extracted from Trachsel et al. (2019) and correspond to a maize experiment comprising 97 genotypes (double haploid) tested in four environments (two reps, and only rep was used for the demo) and scored for grain yield (GY), plant height (PH), anthesis silk interval (ASI), and day to anthesis (DA). Also, genomic (551 marker SNPs) and hyperspectral (five flights - 62 bands per-fly) data were available for analysis. In addition, a kinship matrix was computed using a random sample of SNPs to emulate a pedigree matrix.

